# Microglial synaptic pruning on axon initial segment of dentate granule cells: sexually dimorphic effects on fear response of adult rats subjected to early life stress

**DOI:** 10.1101/2020.10.24.353771

**Authors:** Mario A. Zetter, Angélica Roque, Vito S. Hernández, Oscar R. Hernández-Pérez, María J. Gómora, Silvia Ruiz-Velasco, Lee E. Eiden, Limei Zhang

## Abstract

Axon initial segments (AIS) of dentate granule cells (GC) in hippocampus exhibit prominent spines during early development that are associated with microglial contacts. Here, we asked if developmental changes in axon initial segment spines (AISS) could be modified by neonatal maternal separation through stress hormones and microglial activation and examined the potential behavioral consequences. We examined AISS densities at postnatal day (PND) 15 and 50, using Golgi-Cox staining and anatomical analysis. Neuron-microglial interaction was assessed using antibodies against ankyrinG, PSD95 and Iba1, for AIS, AISS and microglia, respectively, in normally reared and neonatal maternally separated (MS) male and female rats. We observed a higher density of AISS in MS groups at both PND15 and PND50 compared to control. Effects were more pronounced in female than in male rats. AIS-associated microglia showed a hyper-ramified morphology and less co-localization with PSD95 in MS compared to normally reared animals at PND 15. An MS-like alteration in microglial morphology and synaptic pruning could be produced ex vivo by vasopressin application in acute hippocampal slices from normally reared animals. MS rats exhibited increased freezing behavior during auditory fear memory testing which, like effects on AISS density, was more pronounced in females than males. Freezing behavior was associated with Fos expression in dorsal and ventral dentate GC. In summary, AIS associated microglial activity is altered by MS. Sex differences in the long-term effects of MS on AISS density are penetrant to a behavioral phenotype of increased stimulus reactivity in adult female subjects.

## Introduction

Microglia are immune-related resident cells of the central nervous system (CNS) activated for defense against pathogens or injury (Sierra, Paolicelli et al. 2019). More recently, microglia have emerged as a key cell type active in remodeling of brain circuits, including the formation, modification and elimination of synaptic structures, that occurs throughout life span as animals adapt to their environment (Wu, Dissing-Olesen et al. 2015, Sierra, Paolicelli et al. 2019). Microglial cells are produced from yolk sac myeloid progenitors (Ginhoux, Greter et al. 2010) and migrate to the brain around embryonic day (ED) 9.5 in rodents. The immature cells, with amoeboid morphology, migrate through nervous tissue throughout the fetal period, establishing regenerative niches in white matter tracts. As they mature, microglia populate specific brain areas following ontogenic gradients and morphologically differentiate into ramified cells (Morgane S. Thion 2018). By the end of the third postnatal week, microglia attain their adult highly ramified surveillance phenotype. Microglial-mediated synaptic pruning was first demonstrated at the electron microscopy level in 2011 (Paolicelli, Bolasco et al. 2011). Microglia are not a homogenous population of surveilling cells: rather, their density, sensitivity to neurotransmitters or neurohormones, and morphology vary widely across brain regions (Lawson, Perry et al. 1990, Lawson, Perry et al. 1993, Hanisch 2013). For example a microglial population specialized for association with axon initial segment (AIS) of dentate granule cells (DCs) was recently discovered in the dorsal hippocampal formation of rat (Baalman, Marin et al. 2015).

The AIS is defined as a 20-60 μm-long domain located at the proximal axon/soma interface that has a high density of voltage-gated ion channels, at which, the summation of synaptic inputs is believed to give rise to action potentials (Gesell 1940) and for a review, see (Huang and Rasband 2018). The AIS was originally defined ultrastructurally by fasciculated microtubules and an electron-dense undercoat beneath the plasma membrane (Palay, Sotelo et al. 1968). It is defined molecularly by the presence of the master scaffolding protein ankyrinG (Huang and Rasband 2018). Morphological investigations in the late 1970s to early 1980s revealed a significant number of presumably inhibitory synapses on the *shaft* of the AIS of some type of neurons (Somogyi and Hamori 1976, Somogyi 1977, Kosaka and Hama 1979, Kosaka 1980, Somogyi, Freund et al. 1982). These have been implicated in control of neuronal excitability (Bender and Trussell 2012). The presence of AIS *spines* (AISS) in the rat hippocampal formation was also reported at the electron microscopy level in the early 1980 (Kosaka 1980, Kosaka 1983). Moreover, vasopressin (AVP)-containing hypothalamic magnocellular neurosecretory neurons (VPMNN) long-range axon projections were observed to establish Gray type I synapse onto AISS in rat hippocampus (Zhang and Hernandez 2013). However, this phenomenon remains morphologically and functionally underexplored.

Exposure to early life stress (ELS), such as neonatal maternal separation (MS), in altricial species (e. g. human and rodents), is associated with increased fear memory and anxiety-like behavior, such as specific phobia, social anxiety, and panic disorders, that have prominent defensive features (Nelson, Zeanah et al. 2019) (Wang, Levine et al. 2020). The animal model of neonatal MS (Hofer 1973, Plotsky and Meaney 1993), was popularized in the early 90’s by studies showing that extended and repeated periods of maternal separation early in life caused long-term increase in stress reactivity, suggesting that ELS programs animals to be more responsive to stressful situations (Wang, Levine et al. 2020). A key question to be answered is how these changes in behavior are regulated at neuronal circuit level?

The hippocampal formation is a locus for fear encoding, especially for auditory fear learning (Moita, Rosis et al. 2003, Madronal, Delgado-Garcia et al. 2016). It is also one of the most susceptible regions to stress-induced abnormalities in synaptic plasticity (Wei, Simen et al. 2012, Wei, Hao et al. 2015). Abnormalities have been previously observed in hippocampus related to spatial learning in MS subjects (Hernandez, Ruiz-Velazco et al. 2012). We therefore asked if developmental changes in axon initial segment spines, like supernumerary synapse elimination, associated with ELS, might be mediated by microglia activity, under the influence of stress hormones. If so, these could represent a morphological site at which stress acts upon the developing nervous system.

This study aims at evaluating the effects of age, sex and MS on the morphology of GC-AISS, a target region of VPMNNs innervation, which we have demonstrated previously to be potentiated by MS (Zhang, Hernandez et al. 2012). We further sought to correlate these changes with defensive behaviors triggered by an auditory fear memory and immediate early gene c-*fos* protein (Fos) expression in GCs of dorsal and ventral hippocampal formation in adult male and female rats. Our data show that microglial synaptic pruning activity is altered by MS. The long-term effects on synaptic spine density are sexually dimorphic, which may contribute to an altered stress response behavioral profile in adult female compared to male subjects exposed to ELS.

## Materials and methods

### Animals and maternal separation procedure

A total of fifty-one Wistar rats from the local animal facility were used in this study (see Fig. 1 experimental design for details). All animal procedures were approved by the local bioethical and research committees, with the approval ID 085-2013 (renewal 035-2020), in accordance with the principles espoused in the National Institute of Health Guide for the Care and Use of Laboratory Animals (NIH Publications No. 80-23) revised 1996.

**Fig. 1.**
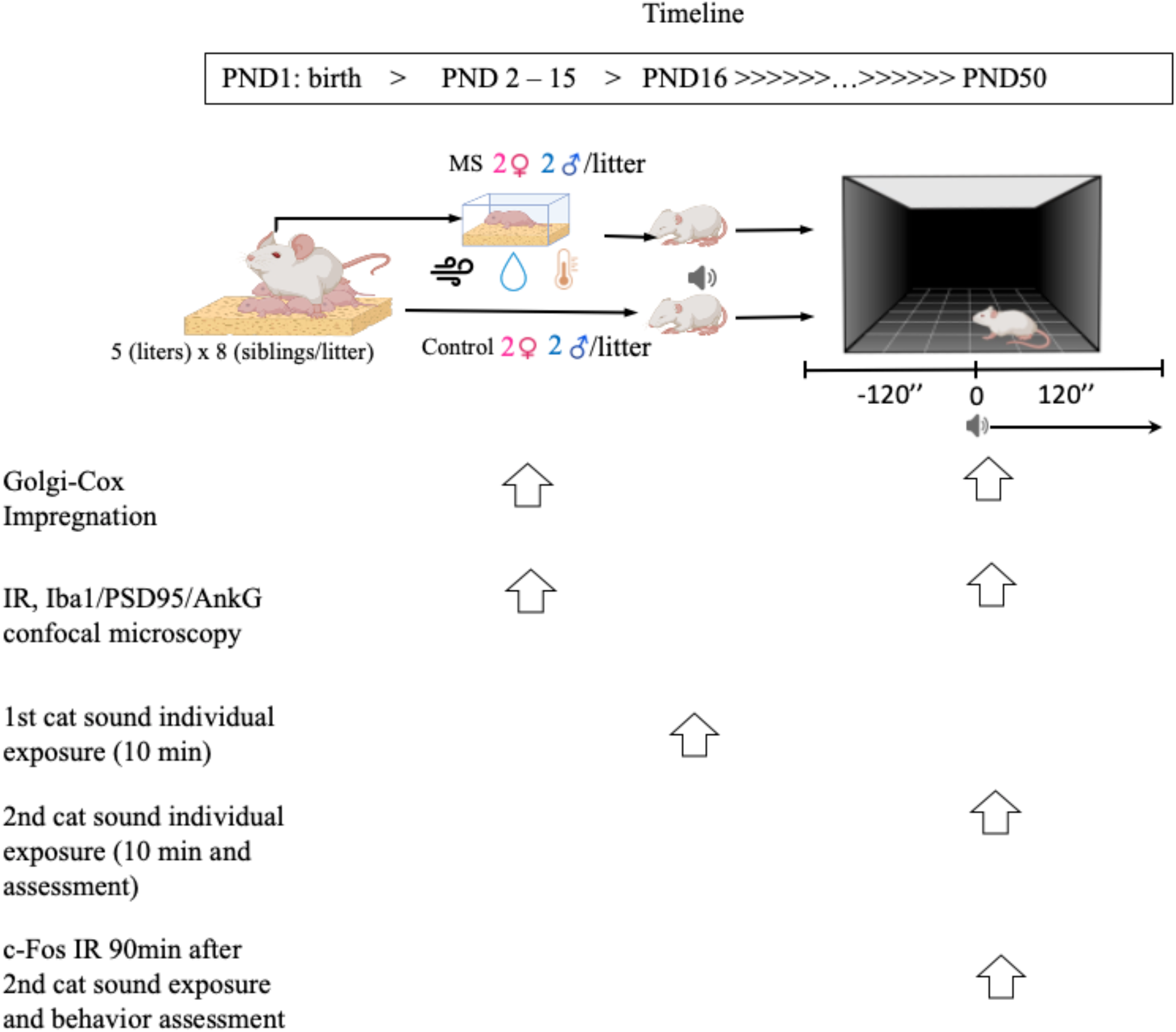
Schematic timeline of the main experimental design. PND: post-natal day; MS: neonatal maternal separation group; IR: immunohistochemical reaction. Symbols: 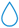 humidity, 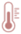 temperature and 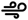 ventilation, 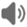 recorded cat vocalization.

The maternal separation (3 h daily) procedure is described elsewhere ((Zhang, Hernandez et al. 2012) with modification according to (Benner, Endo et al. 2014)). Briefly, female and male adult rats were mated for 2 days. During the last week of gestation, female rats were single-housed in standard rat plexiglas cages and maintained under standard laboratory conditions with 12:12 light–dark cycle (light on 0700), temperature maintained at 22 ± 2 °C, and food and water ad libitum. On the day following parturition, PND2, five litters were culled to eight pups, with four males and four females per each litter. During the period from PND2–PND15, half of the pups of each litter were separated from their dams for three hours daily between 9:00 h and 12:00 h. Pups were removed and transferred by gloved hand previously coated with fine bedding-powder from the same cage of each litter. Pups were moved to an adjacent room and placed individually into a small box filled with bedding, and placed in a humid incubator with temperature maintained at 27 ± 1 °C. After the 3 h separation period, the pups were returned to the home-cage with their respective dam. Non-separated siblings were left undisturbed except for changes of the bedding twice a week and served as control groups for this study. Each litter provided n=1 to each experiment, *i.e.,* no more than one sibling within the same liter was an experimental subject.

After PND15 MS procedure, the remaining pups were left undisturbed except for the first predator sound exposure *(vide infra)* and maintained with their dams until weaning at PND28, when males and females were separated and housed in groups of 3 per cage until experimental testing or perfusion.

### Experimental design

Three groups of experiments were performed in this study. Group A were subjects for an integrative assessment of MS effects on development regarding AISS morphometric changes, microglial activity and behavioral consequences concerning fear memory and defensive behavior in adulthood in male and females (see Fig. 1 for detailed experimental design). Five litters with five dams and 40 offspring, 20 male and 20 female rats were used. Group B were subjects for experiments to assess whether or not the stress hormones AVP or PACAP would alter the microglial morphology and PSD95 co-localization, in an ex vivo acute slice pharmacological preparation. Four rats at PND15 that were never removed from the litter were used (two males and two females from animal vivarium). Group C were subjects of experiments in which the morphology of AISS was assessed at a development time point preceding detection of AIS-associated microglia in hippocampus (see (Baalman, Marin et al. 2015)). Two animals at PND5, from animal vivarium, and without removal of pups from litters at any point, were processed to obtain brain tissue for Golgi-Cox impregnation after an overdose of anesthesia and removal of brain after decapitation, without perfusion-fixation (resulted in reduced background staining).

### Golgi-Cox impregnation and morphological analysis of AISS

For Golgi-Cox impregnation procedures, male and female rats of both control and MS groups at PND5, PND15 and PND50 received an overdose of anesthesia and were decapitated (PND5, *vide supra)* or perfused-fixed (PND15 and PND50, *vide infra).* Brains were removed from the skull and were sagittally blocked to make two halves (one for Golgi-Cox impregnation and one for immunohistochemical reaction*, vide infra*). Tissues were briefly rinsed with phosphate buffer 0.1 M (pH 7.4) and then immersed in sequenced impregnation solutions (FD Rapid GolgiStain kit; FD Neuro Technologies, Ellicott City, MD) for two weeks in the dark. Sagittal sections (150 μm thick) were sliced using a vibratome and solution C from the Rapid GolgiStain kit and mounted on gelatin-coated glass slides. These sections were air-dried at room temperature in the dark and then stained with staining solution provided by the kit mentioned previously following provider’s instructions. Samples were examined under microscope and the axon initial segment spines (AISS) of dorsal dentate granule cells were selected to be quantitatively assessed due to its peculiar anatomical feature that allowed us to establish objective criteria for report inclusions (Fig. 2). Counting was made under microscope with experimenter blind to sample groups. Spine density was calculated by tracing a given length of AIS with help of drawing tube (Nikon T550i equipped with a corresponding drawing tube). Finally, representative neurons were selected for 2D-reconstructed of the 3D neuron morphology (150 μm in the Z dimension), using the microscope equipped with a drawing tube.

**Fig. 2.**
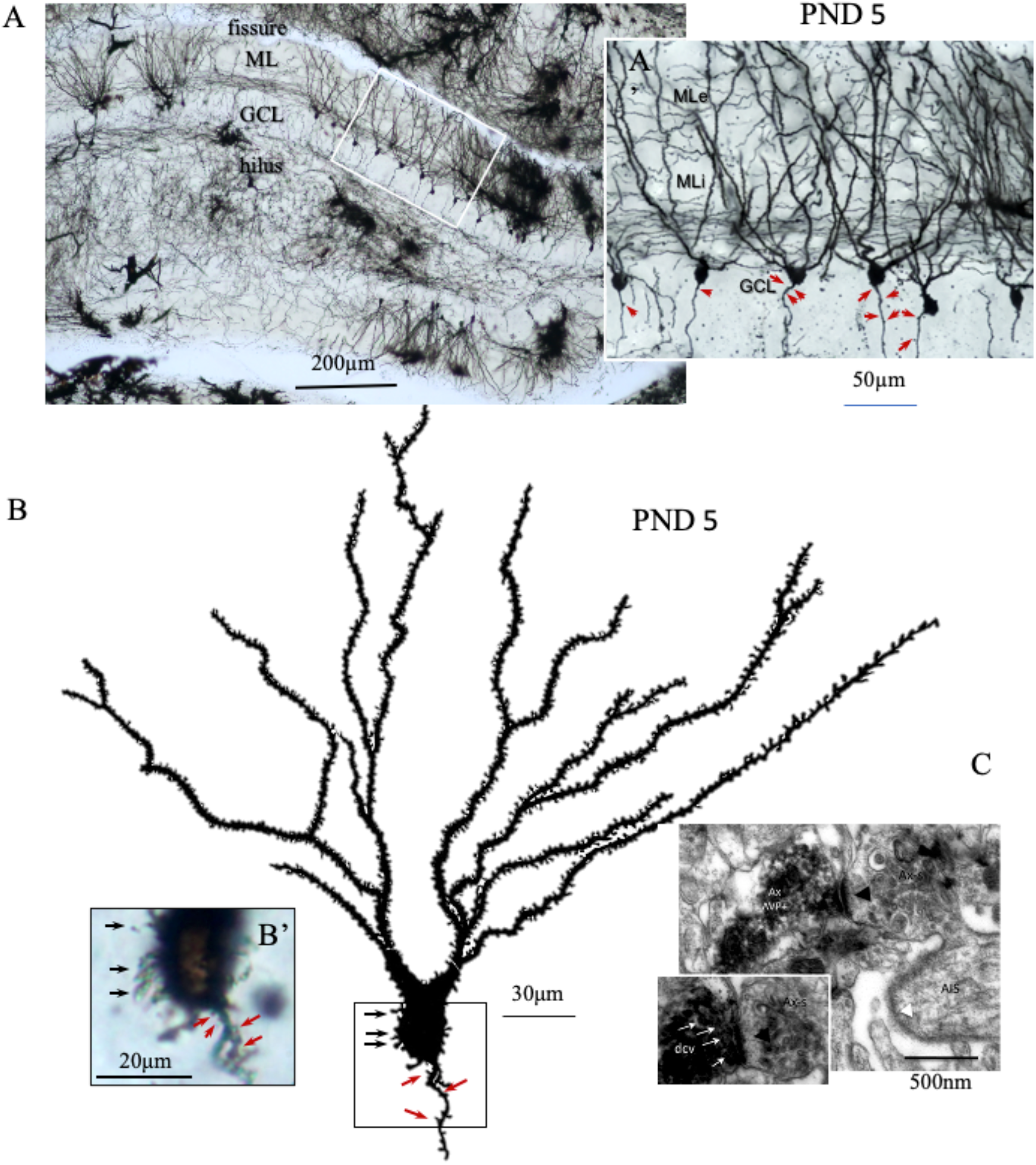
Granule cells (GC) of dentate gyrus (DG) of hippocampal formation displayed abundant somatic and axon initial segment spines during early postnatal stage. **A** and **A’** show numerous GC of DG of a PND5 rat brain, revealed with Golgi-Cox impregnation. Red arrows indicate the spines emitted from the axon initial segment (AISS). **B**. a 2D reconstruction of a 3D (150 μm in Z dimension, the thickness of the brain section) image of a single GC. The somatic spines are indicated with black arrows and the red ones indicate the AISS. **C**. Electron microscopy photomicrograph showing an AISS spine receiving a Gray type I synapse (asymmetric) made by an axon terminal (Ax), which is immunopositive to arginine vasopressin (AVP). White arrows indicate AVP+ dense core vesicles (dcv). AIS: axon initial segment (panel taken from reference (Zhang and Hernandez 2013) with permission).

### Immunocytochemical reactions (IR) for Iba1, PSD95, AnkG and c-Fos

Rats were anesthetized with an overdose of pentobarbital (63 mg/kg, Sedalpharma, Mexico) followed by perfusion/fixation via the transaortic route with NaCl (0.9%) followed by ice-cold fixative (4% paraformaldehyde in sodium phosphate buffer (PB 0.1M, pH 7.4) and saturated picric acid (15% v/v) for 10 min. Brains were removed and thoroughly washed with 0.1 M PB. Free-floating vibratome (Leica VT1000) sections of 70 μm were obtained the day of perfusion-fixation. The following primary antibody cocktails were used: rabbit anti-Iba1 (Biocare, CP 290A, 0.05μg/0.1ml, 1:500, CA, USA); mouse anti-PSD-95 (UC Davis, K2843, 1.0 mg/ml, 1:2000, CA, USA) and mouse anti-ankyrinG (UC Davis/Neuromab 75-146, TC supernatant, 10μg/ml, 1:100). These markers were used to study microglial morphology (Iba1), postsynaptic density protein (PSD-95) engulfment, and axon initial segments (ankG). For immunofluorescence detection, sections were washed and incubated with corresponding fluorochrome-conjugated secondary antibodies: donkey anti-mouse (Alexa 488, Jackson ImmunoResearch, PA, USA) and donkey anti-rabbit (Alexa 594, Jackson ImmunoResearch). We used the expression of the protein product (Fos) of the immediate early gene c-*fos* for neuronal activation assessment. Animals were perfused 90 min after the behavioral experiment (Fig. 1) and IR performed using primary antibody rabbit anti Fos (Santa Cruz, SC-52, 100 μg/ ml, 1:1000, CA, USA) and a biotinylated secondary antibody (goat anti rabbit, Vector, USA), chromogenic development of the reaction was done using the ABC kit (Vector) (for technical details, see (Zhang, Hernandez et al. 2016)).

### Measurements of microglial morphology and PSD95 inclusions

Processing and measurements of microglial cells were made according to Young & Morrison (Young and Morrison 2018). Confocal stack images were obtained from the dorsal hippocampus immunolabeled for Iba1 from P15 rats. Photomicrographs were acquired with an optical section thickness of 1 Airy unit. For the image processing, stacks were flattened into a Z-projection with Fiji (Image J, NIH, MD) and then transformed to 8-bit images. Contrast was adjusted to a pixel radius of 3. Noise was despeckled and the image converted to binary. The image was then skeletonized and trimmed to remove skeleton fragments. The resulting image was analyzed with a Fiji skeleton (2D/3D) plugin and endpoints were summed to obtain a measurement of the number of branches in each microglial cell located in DG from control and MS males and females.

The same imaging parameters were used to obtain photomicrographs of cells immunoassayed for Iba1 and PSD95. In this case, the number of PSD95 inclusions within each microglial cell was counted and the average of puncta in 10 microglia cells (two per rat) was reported for each group.

### Auditory fear memory test and behavioral scoring

We devised a two-step fear memory and defensive behavior test to correlate microglial morphology and AISS changes associated with MS at PND15 with hippocampal neuronal activation, fear memory learning, and defensive behavior in adulthood. The first experimental manipulation was performed on PND16 (Fig. 1), when male and female subjects, both control and MS, were placed individually in a wooden black chamber with dimensions 60×60×40 cm^3^ (Fig. 1), gridded into 36 regions on a cardboard sub-floor. Cat vocalizations (https://www.youtube.com/watch?v=nwv1U7SI-c4) were played for 10 minutes from a speaker over a mesh covering the box. Animals were then returned to their home cages with corresponding dams. The second experimental manipulation was performed on PND50. Animals were individually placed in the ungridded wooden box (Fig. 1), during five minutes for exploration and then exposed to the same sound recording during a second five-minute period. The behavior was video typed. The behavior assessment was made off-line with experimenters blinded to experimental group. The scoring was made every five seconds modified from literature (Detke, Rickels et al. 1995, Zhang, Hernandez et al. 2016, Zhang, Hernandez et al. 2018) during the 120 s before the sound and 120 s after the sound. *Defensive behaviors* were considered as rearing, climbing and freezing, assigning 1, 2 or 3 points respectively, for this study. After the experiment, rats were returned to an isolated sound-insulated location during 90 minutes prior to perfusion-fixation for Fos immunohistochemistry (*vide supra)*.

### Ex-vivo assessment of microglial morphology

Brain slices were prepared from male 15 day-old Wistar rats. Animals were deeply anesthetized with sevoflurane (SevoFlo, Abbott Las, Abbott Park, Illinois) and decapitated. Brains were quickly dissected and sectioned coronally at 300 μm with a vibrating microtome (Leica VT1000 Leica Instruments, Nussloch, Germany). The solutions used for these steps were based on (Weiss and Veh 2011, Zhang, Hernandez et al. 2016). The cutting solution contained (in mM): 75 NaCl, 50 Sucrose, 25 NaHCO3, 2.5 KCl, 1.25 NaH2PO4, 0.1 CaCl2, 6 MgCl2, and 25 D-glucose, pH 7.4. Slices containing dorsal hippocampal formation were incubated in artificial cerebrospinal fluid (ACSF) containing (in mM): 125 NaCl, 25 NaHCO3, 2.5 KCl, 1.25 NaH2PO4, 2 CaCl2, 2 MgCl2, and 25 D-glucose, pH 7.4. Slices were maintained in carbogen bubbling. Slices were divided into three experimental groups: ACSF control, AVP 100nM (kind gift from Professor Maurice Manning) and PACAP 100nM (A1439, Sigma-Aldrich). After the incubation period of 120 min, slices were observed under an inverted (Nikon Eclipse E600FN) microscope with differential interface contrast (DIC) to assure good neuronal viability (small number of picnotic nuclei or swelled cells). Slices were then fixed with PFA 4% for 30 minutes, washed with trizma-base buffer (TBS), and immunoreacted against Iba1 and PSD95, as described above. Finally, slices were mounted with DAPI mounting medium (VectaStain, Vector Laboratories, CA) and the superficial 70μm observed and imaged with a confocal microscope.

### Statistical Analysis

Quantitative results were expressed as mean ±SEM. Groups were tested for normality with a Pearson test. Differences between means were evaluated by three- and two-tailed ANOVA, followed by Tukey post-test. All the analyses were performed using Prism (GraphPad Software, V8.0, San Diego, CA, USA). Differences were considered statistically significant when *p* value was <0.05.

## RESULTS

### AISS decline with development and effect of MS on AISS decline

Using Golgi-Cox impregnation, granule cells (GC) of dentate gyrus of a postnatal day five (PND5) rat exhibited a high occurrence of spines located in soma and axon initial segment (AIS) (Fig. 2 A and B, red arrows). The density of these AIS spines (AISS) was reduced at PND15 (Fig. 3A) and further reduced at PND50 (Fig. 3B). Rats pups that underwent the 3 hr daily maternal separation (MS) during PND-15 exhibited lower AISS reduction than controls. This diminished reduction was more marked in the female group in which, at PND50, the AISS counts were significantly higher than in the male group. In Fig. 3C, we show the result of a three-way ANOVA to test the effects of sex, rearing condition or age on the AIS spine density. Sex effect was not significant (F (1, 152) = 1.3, p=0.26), while age (F (1, 152) = 149, p<0.001) and rearing condition (F (1, 152) = 286, p<0.001) significantly affected the AIS spine density. The three-way interaction (sex*age*rearing condition) was also statistically significant. Tukey’s multiple comparisons test was performed to compare means. The results are shown in Fig. 3C, where columns that do not share any letters have significantly different means. The adjusted *p* values for all the comparisons are shown in table 3D.

**Fig. 3.**
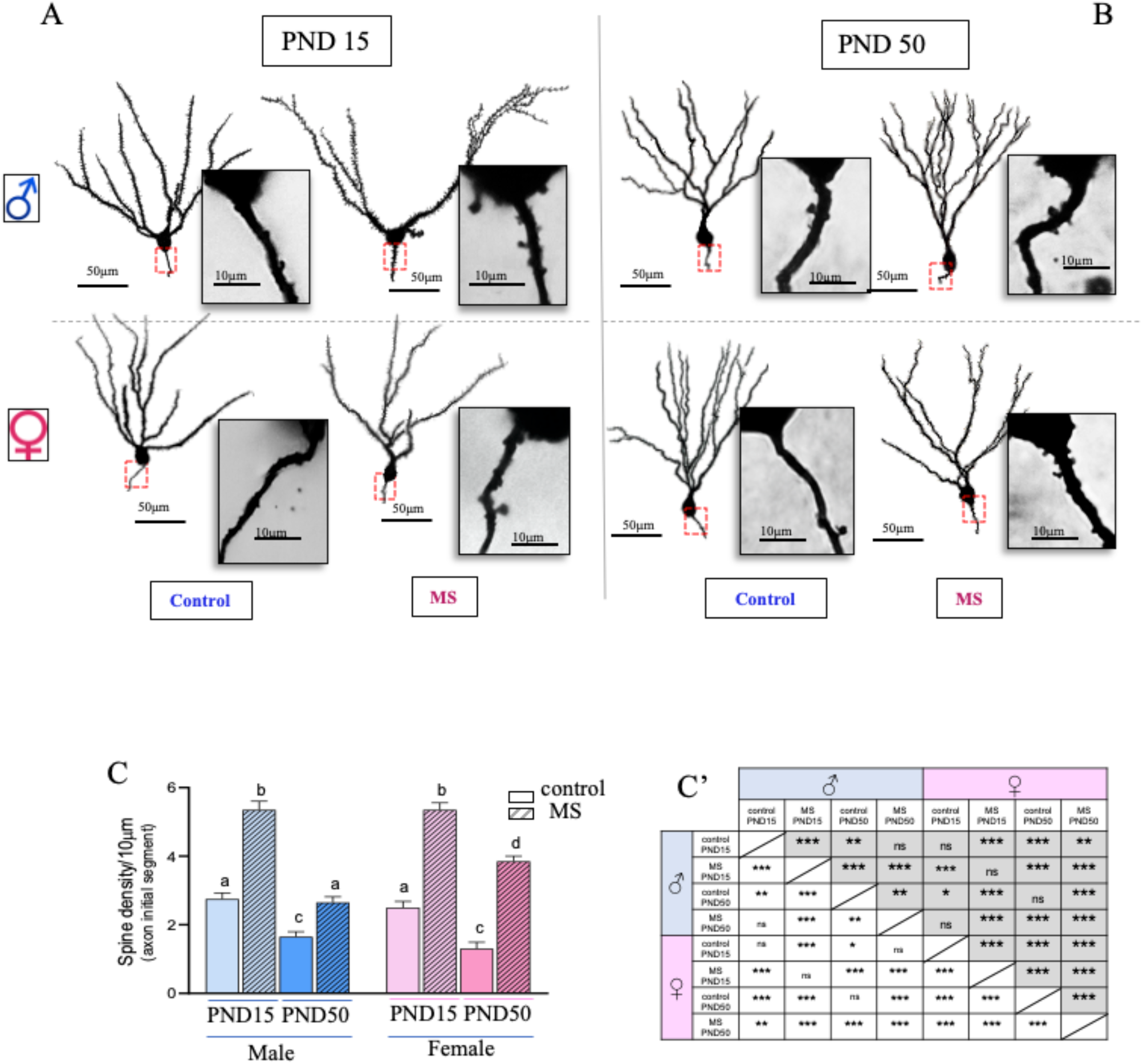
Axon initial segment spines of dentate gyrus granule cells (GC) at both PND15 and PND50 as a function of neonatal maternal separation and sex. A. Camera lucida 2D reconstructions of representative Golgi-Cox impregnated GC (3D with Z=150 μm) and photomicrographs of AIS comparing the AIS spines (AISS) in males and females raised under control or MS conditions at PND15 (**A**) and at PND50 (**B).** In panel (**C**), a three-way ANOVA was performed to see the effect of sex, rearing condition or age on the AISS density. Sex effect was not significant (F (1, 152) = 1.3, p=0.26), while age (F (1, 152) = 149, p<0.001) and rearing condition (F (1, 152) = 286, p<0.001) significantly affected the AISS density, the three way interaction (sex*age*rearing condition) was also statistically significant. Tukey’s multiple comparisons test was performed to compare means, columns that do not share any letters are significantly different, the adjusted *p* values for all the comparisons are shown in table **C’**.

### MS increases freezing behavior and neural activity in the dentate gyrus, with a more pronounced effect in female rats

The dentate gyrus of the hippocampus has been implicated in the maintenance of the episodic and contextual memories (Madronal, Delgado-Garcia et al. 2016, Hainmueller and Bartos 2020). Recent studies have shown its involvement in mapping sounds in relevant behavioral tasks (Aronov, Nevers et al. 2017). We therefore aimed to study if the morphological changes induced by MS at the AISS of dentate gyrus granule cells could be correlated with an increased recall of an aversive auditory stimuli presented at early life and altered behavior in adulthood. To explore this issue, we exposed male and female control and MS subjects to predator sound for 10 minutes when they were at PND15 and evaluated their behavior at PND50 (see Fig 1 for experimental design and Fig. 4 for behavioral results). We found that before exposure to the cat sound, there was no difference in the basal (rearing, climbing or freezing) behaviors displayed by MS or control animals of either sex (front row of Fig. 4A). As expected, exposure to cat sound increased the above-mentioned behaviors in all groups (back row of Fig. 4A). When these behaviors after the cat sound were classified as active (rearing and climbing) or passive (freezing) escaping behaviors (Fig. 4B), it was evident that MS main effect was on the passive behavior (freezing), and that as previously reported, female rats had a higher basal level of passive coping response (Goodwill, Manzano-Nieves et al. 2019) (Papaioannou, Gerozissis et al. 2002). Interestingly MS had a higher impact on freezing in female (*p* < 0.01) than in male rats (*p*<0.05). As AIS are crucial for neuronal depolarization, we examined the state of neuronal activation in dorsal and ventral dentate gyrus (Fig. 4 C and D), subregions that have been typically invoked as having a role in episodic memory / spatial location for the former, and modulation of the stress response for the latter. We found that in both regions, neuronal activation, as measured by Fos immunoreactivity after the behavioral test, was increased in MS compared to control rats, and accordingly, that the differences were greater between female MS and control rats than between male MS and control rats (Fig 4 E and F).

**Fig 4:**
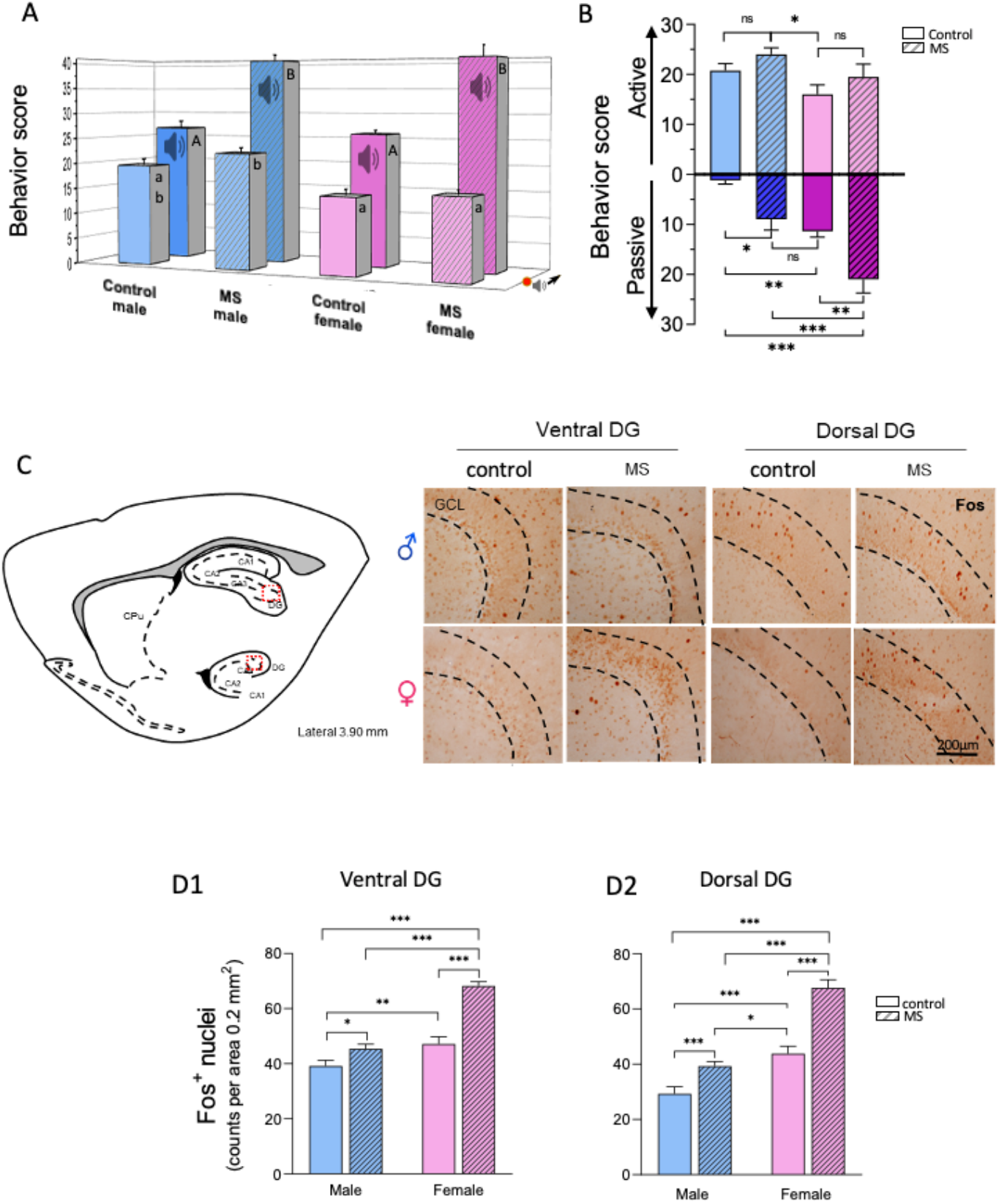
Maternal separation differentially impacts differentially fear memory GC neuronal activation and defensive behavior in young adulthood in male and female offspring. **A:** open field box test performed at PND50, with 2nd predator sound exposure (first was at PND15) triggered *defensive behavior* in rats (scoring points assignation: freezing=3 points, climbing = 2 points, and rearing = 1 point, n=5 per group). The bars in the front row were scores obtaining analyzing the 120 seconds prior the predator sound started (symbolized by 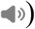. A two-way repeated-measures ANOVA with “sound exposure” and “experimental group” as factors showed a significant effect of sound exposure (F (1, 16) = 245 *p*<0.001), experimental group (F (3, 16) = 30 *p*<0.001) and interaction (F (3, 16) = 16 *p*<0.001). Tukey’s *post hoc* test was used to evaluate differences between experimental groups before and after cat sound exposure. Columns with different letters indicate significant differences of means, lowercase letters were used to compare groups’ behavior before cat sound (front row) and uppercase letters were used to compare groups’ behavior during cat sound. There was a significantly higher score in the defensive behaviors of maternally separated rats (MS male 40.60 ± 1.33 and MS female 41.00 ± 2.35) compared to animal facility reared rats (control male 26.80 ± 1.39 and control female 26.00 ± 0.7) during the cat sound exposure. **B:** *Defensive strategy* was itemized as *active* (climbing and rearing, upright bars) and *passive* (freezing, inverted bars), during cat sound exposure. A two-way ANOVA using sex and rearing condition as factors showed that, for the active strategy there was a significant effect of sex F (1, 16) = 7.527, *p*=0.01 but not for rearing conditions or for the interaction between factors. Tukey *post hoc* test showed significant differences only between male MS (24 ± 1.34) and female control (16 ± 1.92) groups. For the passive strategy, a two-way ANOVA showed a significant effect of sex F (1, 16) = 45.63, *p*<0.001; rearing condition F (1, 16) = 28.03, *p*<0.001; and no interaction between factors. Tukey *post hoc* test showed significantly higher scores in MS compared to control rats (male control: 1.2 ± 0.7 vs male MS: 9 ± 2.1, p< 0.05 and female control: 11.4 ± 1.12 vs female MS: 21 ± 2.1, p<0.01). Also, female rats showed higher scores compared to male rats in the same rearing conditions. Asterisks depict significant differences with values as follows: * *p*<0.05; ** *p*< 0.01; \*\*\**p*< 0.001. **C:** Neuronal activation in the ventral and dorsal dentate gyrus was analyzed by counting the number of Fos+ nuclei in an area of 0.2 μm^2^ along the cell body layer. The drawing shows the lateral coordinates and regions where Fos+ nuclei were counted, and the photomicrographs show examples of immunoreactivity against Fos in the ventral and dorsal dentate gyrus (DG) in control and MS rats of both sexes. A two-way ANOVA was run on a sample of n = 10 observations per group, to examine the effect of sex and rearing condition on the number of Fos+ nuclei in ventral (D1) and dorsal (D2) dentate gyrus. There were significant effects of sex (ventral DG: F (1, 36) = 143, p<0.001 and dorsal DG: F (1, 36) = 187.0, *p*<0.001), rearing (ventral DG: F (1, 36) = 115, *p*<0.001 and dorsal DG: F (1, 36) = 88.87, *p*<0.001) and interaction (ventral DG: F (1, 36) = 38.8, *p*<0.001 and dorsal DG: F (1, 36) = 11.84, *p*<0.001) on the number of Fos+ nuclei. *Post hoc* Tukey test in the ventral DG showed a significantly higher number of Fos+ nuclei in MS groups compared to control in males (control: 39.2 ± 1.42 vs. MS: 45.4 ± 1.16) and females (control: 47.1 ± 1.8 vs. MS: 70.5 ± 0.97). In the dorsal DG significantly higher Fos+ nuclei were found in MS compared to control rats in males (control: 29.2 ± 1.73 vs. MS: 39.2 ± 1.14) and females (control: 46.3 ± 1.73 vs. MS: 67.8 ± 1.97).

**Fig 5:**
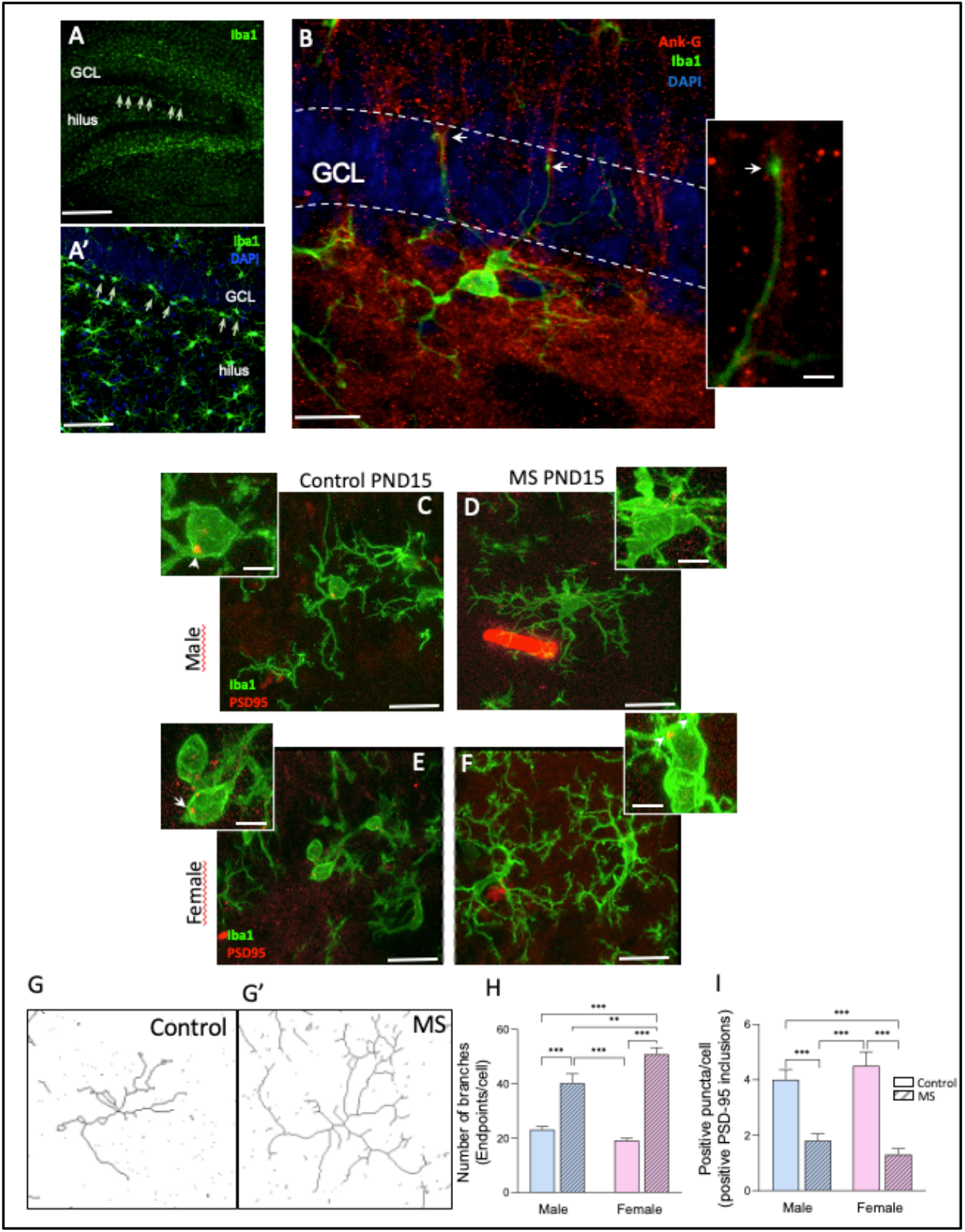
MS triggered hyper-ramified morphology of AIS associated microglia and decreased engulfment of PSD-95 puncta. **A and A’:** a subpopulation of Iba1 immunoreactive (IR) cells (microglia) was observed to be alienated in the sub-granule cell layer of the dentate gyrus (indicated by white arrows). **B.** Microglial Iba1 positive processes were observed running in parallel, in close proximity to ankyrinG immunopositive AIS of the granule cells. Arrows indicate the contact points of microglial process (Iba1 IR). **C-F:** Photomicrographs showing examples of the changed microglial (Iba1 IR) morphology (hyper-ramified in the group of MS) and PSD-95 positive *puncta* located inside microglial cells at subgranular zone of DG. **G and G’:** Representative skeletonized microglial cells of control and MS rats (see method section for details). H: Two-way ANOVA analysis of number of branching in different groups showed that even that the effect for sex is not significant, it was for rearing condition (F (1, 36) = 120.7*, p*<0.0001) and the interaction (F (1, 36) = 10.99, *p*=0.0021). Post hoc Tukey test showed significantly higher number of endpoints in MS rats in males (control: 23.12 ± 1.1 vs MS: 40.2±3.49, *p*<0.001) and females (control: 19.06 ± 0.82 vs. MS: 50.9 ± 2.39, *p*<0.001). I: analysis of PSD-95 puncta within Iba1 labeled cells in different groups showed no effect for sex or interaction, therefore we performed a t test to find the difference of rearing: male (control 4 ± 0.36 vs. male MS: 10.8±.25, *p*<0.001) and female (control: 4.5 ± 0.5 vs. female MS: 1.3 ± 2.1, p<0.001). ** *p*< 0.01; *** *p*< 0.001.

### Microglial morphology and synaptic pruning at dentate gyrus are altered by MS

Since microglia cells have been associated with synaptic pruning we evaluated if there were changes in microglial morphology associated with MS. Microglia display a wide array of morphological adaptations depending on their activity and physiological state, ranging from an amoeboid-highly phagocytic morphology to a ramified, surveyor one (Fernández-Arjona, Grondona et al. 2017). As microglia mature, they transit from an amoeboid to a ramified morphology (Gómez-González and Escobar 2010). This functional plasticity allows microglia to establish an exceptional immune and surveying network in the brain. Widely different distribution profiles exist between microglia depending on their localization in the CNS (Jinno, Fleischer et al. 2007), and a specific subclass of microglia associated to axon initial segments has been recently described (Baalman, Marin et al. 2015). Using Iba1 immunofluorescence staining we noted a group of microglial cells located in the subgranular cell zone (sGCZ) of the dentate gyrus, facing the *hilus,* where the axons (mossy fibers) of granule cells leave the GCL (Fig 4 A and A’). Staining with an antibody against ankyrinG, a scaffold protein located at axon initial segments, revealed that the processes of the microglial cells in the sGCZ were contacting axon initial segments located in the inner layers of the GCZ (Fig. 4 B, and inset). Then, we asked whether microglial in the sGCZ might be differentially associated with the pruning of excitatory synapses in control or MS animals. With dual immunofluorescence for Iba1 and PSD-95 at PND15, we found that MS microglial cells showed a hyper-ramified (surveying) morphology that was associated with less colocalization with PSD-95 positive puncta in the cytoplasm of Iba1 positive cells while control microglia showed an amoeboid morphology and more PSD-95 puncta in their cytoplasm (Fig 4 C-I), suggesting that MS inhibited the normal engulfment of postsynaptic excitatory spines in the AIS of granule cells.

### The stress-related peptides vasopressin and PACAP regulate the morphological and functional changes observed in MS microglia

Previously, we have demonstrated that magnocellular vasopressinergic neurons of the hypothalamus synaptically innervate the dorsal and ventral hippocampus, (Zhang and Hernandez 2013) and the presence of vasopressin immunopositive synaptic terminalis making synaptic contacts with AIS (Fig. 2 C). Also, we have shown that maternal separation potentiates the vasopressinergic system in the paraventricular and supraoptic nucleus, which is correlated with increased vasopressinergic innervation of the hippocampus and amygdala (Zhang, Hernandez et al. 2012, Hernandez, Hernandez et al. 2016) and that the hippocampus is importantly influenced by other regulatory peptides such as PACAP and its receptor PAC1, whose mRNAs is abundantly expressed in hippocampus (Zhang, Hernández et al. 2020).

Accordingly, we have asked whether neuropeptides such as vasopressin or PACAP - with known regulatory effects on microglia (Ringer, Buning et al. 2013), might contribute to the morphological findings in MS animals. In concordance with the hypothesis that regulatory peptides might drive the shape of the adult structure of granule cells, by regulating the pruning activity of microglia, we found that incubation of acute brain slices with vasopressin (100 nM) during two hours resulted in a hyperramified microglial morphology with scarce, if not absent colocalization of PSD-95 *puncta* in comparison (Fig. 6A) with slices incubated with ACSF in which microglia had a more retracted morphology and abundant PSD-95 and Iba1 colocalization (Fig. 6B). In contrast, incubation with PACAP 100 nM induced an alternative, bushy morphology with large swollen somata and short branches, that has been related to increased migratory and phagocytic activity of microglia. Abundant PSD-95 positive *puncta* located in the cytoplasm of these cells suggests that PACAP stimulates spine phagocytosis, while vasopressin exerts an inhibitory role in the spine phagocytic function and the pruning activity of microglia.

**Fig. 6.**
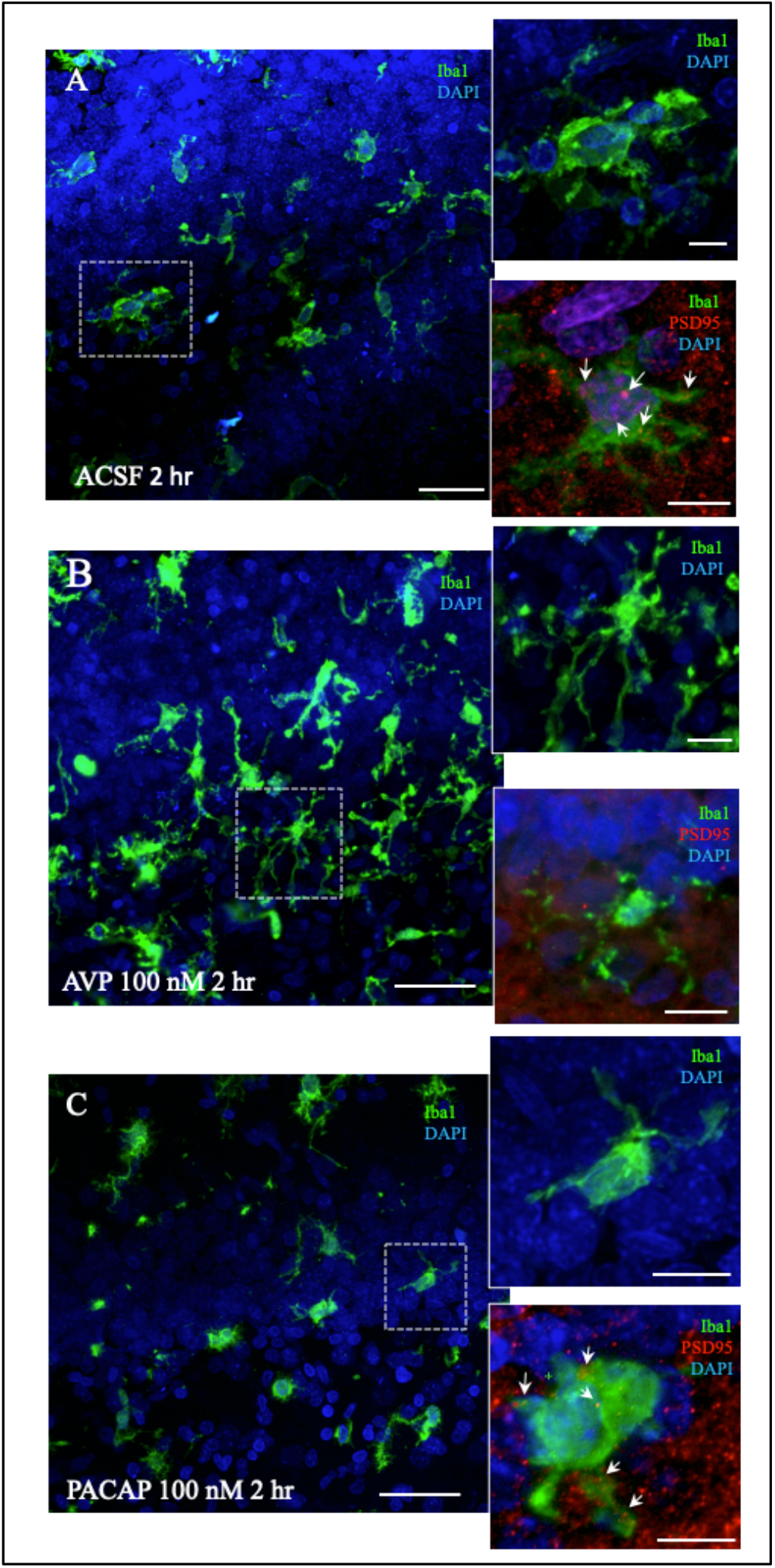
Pharmacological treatment in ex-vivo hippocampal slices with AVP mimicked the morphological findings in MS while PACAP treatment reduced microglial branching. **(A)** AIS associated microglia in acute hippocampal slice treated with artificial cerebral-spinal fluid (ACSF). Insets show the Iba1-IR cells morphology and numerous PSD-95 labeled puncta (engulfed spine remains) within the microglia cells. **B.** treated with arginine vasopressin 100 nM in ACSF for 2hrs. Insets show the hyper-ramified Iba1IR cell morphology and lack of PSD-95 puncta inside the microglia cell. C. treated with PACAP 100nM in ACSF for 2hrs. Insets show the bushy, phagocytic-associated morphology, with large soma and short branches of Iba1IR cell morphology and abundant colocalization of PSD-95 *puncta* inside the microglia cell.

## Discussion

In this study, we investigated the effects of ELS, in the form of maternal separation, in the morphology of granule cells of dentate gyrus of hippocampus, a target of stress-related neuropeptides. The morphological analysis of granule cells revealed that during early life in male and female rats, the axon initial segments feature abundant spine-like processes (AISS) which are reduced at PND50. We found that MS interferes with this physiological reduction of AISS in both sexes, but to a greater extent in females.

Hippocampus is a well-studied brain area related to the cognitive and behavioral responses to stress and fear. In particular, function of the dentate gyrus granule cells of the hippocampal formation has been related to the recall of fear. We therefore attempt to determine whether AISS reduction might be associated with associative memory recall of an aversive stimuli received in early life. We found that male and female MS rats responded more strongly to predator sound than controls. Additionally, when active and passive responses were separately assessed, MS females responded to fear with marked freezing (passive) behavior. This may represent a more complex behavioral response, and could reflect a greater susceptibility to anxiety, a more efficient survival response, or indeed both de Kloet and colleagues have proposed that the switch from active to passive coping strategies in the forced swimming test is not necessarily a signal of depression but an appropriate survival-oriented learning mechanism (Molendijk and de Kloet 2015, de Kloet and Molendijk 2016).

The long lasting modifications in the AISS induced by a stressful situation in early life, resonate with the classical notion that memory lies within the strength of the synapse (Cajal 1894, Hebb 2002). Under this framework, we hypothesize that, in response to ELS, microglia could be a bridge between external stimuli and synaptic modification, whereby environmental cues sculpt the brain at the circuit level. First, we observed that the anatomical distribution of microglial cells near to subgranular cell zone (sGCZ), where the axon initial segments of granule cells are, could be a key area for modulating the hippocampal circuit activation, as GC are the main relay of inputs from cortical information to the hippocampal formation. This organized localization suggests that some ontogenically-derived mechanisms attract the highly motile microglia to this area early in development (before PND15), probably to survey and prune the AISS of granule cells. In fact, in other brain areas as the visual system and the auditory cortex, microglia operate upon the supernumerary synaptic spines in an activity-dependent manner in a such way that the absence of visual or auditory stimuli disrupt this process and lead to abnormalities in synaptic transmission in these areas (Paolicelli, Bolasco et al. 2011, Squarzoni, Oller et al. 2014, Milinkeviciute, Henningfield et al. 2019).

The distribution of microglial cells, together with the localization of their branches near ankyrinG-positive AIS, supports recent observations which point to a subclass of microglial cells associated to AIS that migrate during early development (PND7-9) and associate mainly with excitatory neurons (Baalman, Marin et al. 2015). AIS are crucial neuronal segments, as the axon potential initiates here, due to its lack of myelin and the abundant presence of sodium and calcium voltage dependent channels (Colbert and Johnston 1996, Bender and Trussell 2009). Thus, the presence of excitatory synapses in this neuronal region could alter the neuronal compartmentalization and result in neuronal hyperexcitability.

Previous studies have shown that humans exposed to ELS have increased connectivity at amygdala in association with individual hyperreactivity (Johnson, Delpech et al. 2018). This is in concordance with our results showing that normal pruning of granule cells AISS by microglia is impaired by maternal separation. The maintenance of AISS in granule cells might result in an overactivation of the functional output of dorsal hippocampal circuits related to memory and place location and ventral hippocampal circuits related to modulation of the stress response. Hence, the effects stress during early life on the phagocytic activity of microglia might be a critical link in configuring a circuit that is adaptatively tuned to be more alert in a hostile environment. A previous study where microglia were depleted in early development by central infusion of liposomally encapsulated clodronate, showed that individuals exhibited changes in social locomotor and mood related behaviors, with differential penetrance in males and females (Nelson and Lenz 2017). Most notably decreased ‘behavioral despair’ in the forced swim test after interruption of normal microglial function is reminiscent of the effects on passive and active stress coping reported here.

With the demonstration of a critical microglial link between environmental activity and synaptic plasticity, how might external environmental inputs related to stress, threat and survival be conveyed to microglial hippocampal cells? One mechanism is suggested by selective innervation of limbic structures, including the hippocampus, by neuropeptides within circuits highly associated with the stress response. As an example, we have previously reported that in response to ELS, vasopressinergic system is upregulated and that axonal projections of VP MMNs synaptically connect with AIS spines, establishing symmetrical (excitatory) contacts in all fields of hippocampal formation, these synaptic boutons contain large dense core vesicles (LDCV) which are known to contain neuropeptides as AVP and notably innervate DG granule cells AIS (Zhang, Hernandez et al. 2012). Thus, the upregulation of this stress-related system by MS and the increased release of AVP to the microenvironment of AIS might induce the microglia to acquire a passive phenotype and to reduce the pruning activity.

Surprisingly, to the best of our knowledge, no reports in literature show information about the role of neuropeptides in the regulation of synaptic pruning in hippocampus; however, we consider this point crucial for understanding the circuit refinement in response to environmental cues as different peptidergic signals could differentially affect microglial activity. For instance *Pituitary Adenylate Cyclase Activating Polypeptide* (PACAP) exert modulatory effects on microglia through acting in PAC1 receptors (Wainwright, Xin et al. 2008). In turn, microglia increase the production of transforming growth factor beta (Carniglia, Ramirez et al. 2017), which has been associated to the stimulation of synaptic pruning (Bialas and Stevens 2013). In the case of vasopressin, only few reports have assessed the AVP receptor expression and effects on microglial function (Pannell, Szulzewsky et al. 2014).

Our results with acute brain slices incubated with regulatory peptides strongly suggest that these could be important factors regulating synaptic pruning and thus play a role as a bridge between neurons (somatosensory inputs) and microglia (morphological sculptors of neural circuits), to “translate” the information from the environment to the structure and synaptic plasticity of neural circuits. In accordance with this hypothesis, we found that incubation with AVP evoked the hyper-ramified phenotype, which was associated with less PSD-95 positive *puncta* in the cytoplasm of Iba1 positive cells, an scaffolding protein located in active excitatory synapses used to indicate spine phagocytosis in physiological and pathological conditions (Paolicelli, Bolasco et al. 2011). In the other hand, the treatment of hippocampal slices with PACAP induced morphological features that characterize bushy / hypertrophic microglia with big soma and short branches, which has been associated with phagocytic activity and increased migratory state (Karperien, Ahammer et al. 2013). Moreover, the PSD-95 IR localization in cytoplasm of these cells was increased. This observation suggests that neuropeptides such as AVP and PACAP modulate the surveillant/phagocytic function of microglia in activity dependent manner, and that the exposure to ELS influenced the molecular pathways that govern the microglial-neuron interactions resulting in a response oriented towards maintaining, rather than eliminating, synaptic sites. It will be an urgent research priority in testing this hypothesis to directly assess the impact of neuropeptide dysregulation, pharmacologically and/or genetically, on MS-mediated synaptic plasticity and consequent changes in adult behavior, and to explore in such models’ cellular changes in hippocampal microglia that may link the two.

MS elicited a dimorphic effect, that was more pronounced in females than males, whose granule cells at DG featured more AISS and neuronal activation in response to environmental threatening cues. MS females responded with more freezing behavior, suggesting that ELS elicited an adaptive response that primed neural circuits to fast respond, and thus, this could be interpreted as a more successful strategy for avoiding threatens to their current environment. This is in accordance with previous studies that revealed that, in threatening conditions, males explore and feature a more risk-prone behavior than females, which show aversive behavior towards threatening conditions (Jolles, Boogert et al. 2015). The presence of AISS by the exposure to ELS could exacerbate these differences priming individuals for more survival chances when exposed to environmental threats. One possible mechanistical explanations for this sexually dimorphic effect is that sex influences microglial function. In the developing brain, chromosomal sex, gonadal hormone exposure or differential microenvironmental signals might play a role. However, since microglia expression at early postnatal age is very low the most likely mechanism is that prenatal hormone milieu induces differential changes in the microenvironment of males and females and is this different microenvironment the responsible for differences in the activity of microglial activity (VanRyzin, Marquardt et al. 2020). For instance, It has been demonstrated that the perinatal surge of testosterone induces an elevated cannabinoid tone in amygdala and this in turn induces a phagocytic ameboid phenotype in male microglia (VanRyzin, Marquardt et al. 2019). To the best of our knowledge, a similar sex-dependent differential activity of microglia induced by different microenvironment has not yet been yet reported in hippocampus. However it is known that the vasopressinergic innervation of hippocampus is highly dimorphic (De Vries and al-Shamma 1990). As for the exploration of neuropeptide modulation in the context of ELS-mediated changes in hippocampal plasticity and adult behavior, these inquiries will require more sophisticated molecular and cellular investigations in both rats and mice.

## Author Contributions

MAZ, AR and LZ conceived the study and discussed with all authors. MAZ, AR, VSH, ORHP performed the animal experiments. VSH, MAZ, AR, MJG and LZ performed microscopical studies. VSH, SRV, MAZ, LEE and LZ performed statistics analysis. MAZ, VSH and LZ prepared the figures. LZ and LEE provided research resources. MAZ, VSH, LZ, LEE wrote the initial manuscript. All authors edited and discussed.

## Acknowledgements

We thank Mr. Manuel Hernández for producing the modified open-field box for the predator sound test, to Dr. Rafael Hernández for providing animal facility assistance. MAZ thanks CONACYT postdoctoral fellowship through CONACYT-CB–238744. AR is a doctoral student from *Programa de Doctorado en Ciencias Biomédicas, Universidad Nacional Autónoma de México (UNAM)* and was supported by CONACYT (CVU 509694/fellowship No. 286252). OH-P is a postdoctoral fellowship *from Programa de Becas Posdoctorales* of DGAPA/UNAM. This study was supported by grants: UNAM-DGAPA-PAPIIT-IN216918 & CONACYT-CB–238744.

